# Transition paths across the EMT landscape are dictated by network logic

**DOI:** 10.1101/2024.12.03.626660

**Authors:** Anupam Dey, Adam L. MacLean

**Author notes:** Correspondence (A.L.M.).

## Abstract

During development and cancer metastasis, cells transition reversibly from epithelial to mesenchymal via intermediate cell states during epithelial-mesenchymal transition (EMT). EMT is controlled by gene regulatory networks (GRNs) and can be described by a three-node GRN that permits tristable EMT landscapes. In this GRN, multiple inputs regulate the transcription factor ZEB that induces EMT. It is unknown how to choose the network logic for such regulation. Here we explore the effects of network logic on a tristable EMT network. We discover that the choice of additive vs multiplicative logic affects EMT phenotypes, leading to opposing predictions regarding the factors controlling EMT transition paths. We show that strong inhibition of miR-200 destabilizes the epithelial state and initiates EMT for multiplicative (AND) but not additive (OR) logic, suggesting that AND logic is in better agreement with experimental measurements of the effects of miR-200 regulation on EMT. Using experimental single-cell data, stochastic simulations, and perturbation analysis, we demonstrate how our results can be used to design experiments to infer the network logic of an EMT GRN in live cells. Our results explain how the manipulation of molecular interactions can stabilize or destabilize EMT hybrid states, of relevance during cancer progression and metastasis. More generally, we highlight the importance of the choice of network logic in GRN models in the presence of biological noise and multistability.

## Introduction

The fate of a cell is determined by its transcriptional state, which is in turn determined by a gene regulatory network (GRN). GRNs consist of signed (activating or inhibitory) directed interactions between transcription factors, genes, and other genetic elements (Britten et al., 1969). Constructing “complete” GRNs that describe biological function is typically beyond our grasp. With a few possible exceptions such as sea urchin development (Davidson et al., 2002) or partial mammalian networks, e.g. in hematopoiesis (Schütte et al., 2016), constructing whole-organ GRNs remains a grand challenge. Moreover, GRNs quickly grow large and contain many redundancies (Iwasa, 2023). Rather than seeking to construct full networks, small GRNs (i.e. 2-5 node) network “motifs” (Alon, 2020) can explain expected phenotypes with low-dimensional dynamics (Tripathi et al., 2023). Geometrical models offer means to explain low-dimensional dynamics by providing a bridge between the metaphorically smooth Waddington landscape (Waddington, 1957) and quantitative measurements of single-cell states (Rand et al., 2021; Sáez et al., 2022). A common challenge in the construction of models described by small network motifs is the choice of network logic.

Epithelial-mesenchymal transition (EMT) is a quintessential cell state transition, and can be characterized by small GRNs (Hong and Xing, 2024). Various GRN models of EMT have been proposed, most of which share a core network comprised of transcription factors ZEB and SNAIL, and micro-RNAs miR-200 and miR-34 (Hong, Watanabe, et al., 2015; Jolly et al., 2016; Lu et al., 2013; Tian et al., 2013; Zhang, Tian, et al., 2014). The core network consists of mutual inhibitory feedback loops between ZEB—miR-200 and SNAIL—miR-34. Expression of miR-200 maintains the epithelial state, while ZEB (representing the transcription factors ZEB1 and ZEB2) induces EMT. ZEB can be activated by SNAIL, which itself is inhibited by miR-34. Models have shed light on the dynamics and the cell states accessible during EMT: in particular, at least three stable steady states can exist during EMT and cells often transition through one or more EMT intermediate cell states. EMT intermediate states are also referred to as partial EMT states or hybrid E/M states (Sha et al., 2019). In addition, EMT can proceed through multiple different transition paths (Wang et al., 2022).

To construct a mathematical model from a GRN (described by ordinary or stochastic differential equations) a choice of network logic must be made when a gene receives more than one regulatory input. The model of Tian et al. (2013), for example, is constructed with additive gene regulation (OR logic) whereas the model of Lu et al. (2013) is multiplicative (AND logic). There is in general no principled guidance on the choice of network logic when constructing a mathematical model. Indeed, recent work has shown that signals combine to regulate gene expression in both additive and multiplicative manners at roughly equal proportions (a variety of other responses are also observed at lower frequency) (Sanford et al., 2020). The gene-level specificity of the responses measured by Sanford et al. (2020) (for just two specific input signals: retinoic acid and TGF-*β*) emphasizes the challenge in choosing network logic to model a GRN. Furthermore, the impact of this choice on gene regulatory dynamics and the resulting cell fate landscape is for the most part unknown.

In this work we investigate how the choice of network logic impacts gene regulatory dynamics and cell state transitions during EMT. We do so via a GRN that permits tristable EMT, consisting of SNAIL, miR-200, and ZEB (Jia, Jolly, Tripathi, et al., 2017; Lu et al., 2013). We discover that EMT phenotypes are sensitive to the choice of logic. In the next section we introduce the GRN model and discuss the transition paths it permits. We go on to show that the EMT regulatory parameters have opposing effects on the landscape depending on the choice of logic. We first demonstrate this for a constrained model with comparable interaction strengths, and then show that our results hold for a wide range of models with unconstrained parameters. We discuss how the combinatorics of gene regulation can explain the opposing effects observed. Finally, we show how GRN logic can be inferred from experimental data in cells undergoing EMT.

## Results

### A three-node network characterizes cell states accessible during EMT

To investigate how the direct and indirect regulation of ZEB impacts the tristable dynamics of EMT, we studied a three-node gene regulatory network (GRN), introduced by Lu et al. (2013). The GRN consists of SNAIL (*S*) which acts as an input signal, miR-200 (*A*), and ZEB (*B*). *A* and *B* mutually inhibit each other. *S* inhibits *A* and activates *B*; and *B* also has a self-activation (Fig. 1A). Of note: in this network *B* can be both directly activated by *S* and indirectly activated by *S* via a double inhibition: *S* ⊣1 *A* ⊣1 *B*. The direct and indirect activation pathways form a coherent feedforward loop (Alon, 2020) that regulates *B* in combination with two additional loops: mutual inhibition between *A* and *B* and self-activation of *B*. Modeled by ordinary differential equations (ODEs) with Hill function kinetics, this model permits bistability or tristability under certain conditions.

**Figure 1:**
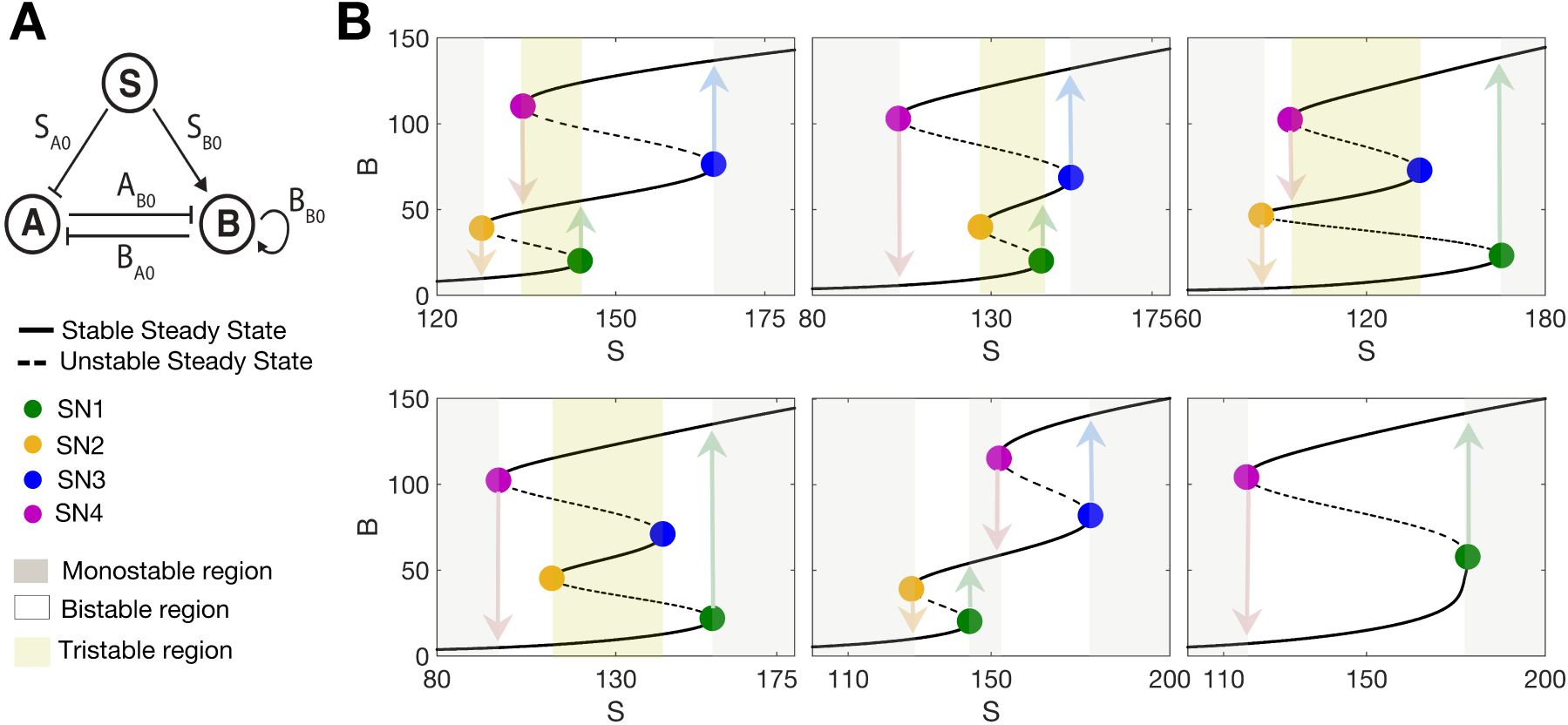
Multistable responses generated by a three-node EMT network. **A**. The three-node EMT network described by input signal SNAIL (*S*), miR-200 (*A*) and ZEB (*B*). Parameters (*S_A_*_0_*, S_B_*_0_*, A_B_*_0_*, B_A_*_0_*, B_B_*_0_) denote the half-maximal concentrations of species that (positively or negatively) regulate another species in the network. **B**. All possible multistable responses in *B* with respect to *S*. Four types of tristable response and two bistable responses exist; colored circles denote saddle node (SN) points. Number of upward transitions marked as 1U or 2U; similar for downward transitions (D).

To construct an ODE model for a GRN, one must choose how multiple inputs regulating a gene affect its expression. AND logic describes responses that are multiplicative with respect to multiple inputs; OR logic describes responses that are additive with respect to multiple inputs There is little principled guidance in the literature regarding this choice, as highlighted by comparison of two well-studied models of EMT in the literature. Lu et al. (2013) model the GRN with AND logic whereas Tian et al. (2013) model the GRN with OR logic. Appealing to experimental evidence yields little guidance: when two signals were combined experimentally to study the logic of gene expression responses, both additive and multiplicative responses were observed in almost equal proportion (as well as less common sub-additive or super-multiplicative responses) in an interaction-specific manner (Sanford et al., 2020). Thus, it is difficult to make principled guesses a priori, and to be confident in any particular choice would require measurement of specific gene-gene interactions in response to a specific stimulus. To account for this uncertainty, we consider models of the same GRN constructed with alternative logic, with the AND model is given by Eqns. 1 and the OR model is given by Eqns. 2, to investigate how network logic impacts the cell states that are accessible during EMT.

The minimal network that permits tristability consists of two coupled positive feedback loops with ultrasensitivity and can be found in EMT as well as other biological contexts (Dey et al., 2021; Frankhouser et al., 2024). The three-node EMT network considered here permits tristability for both AND and OR logic, displaying six different multistable responses in *B* to changes in the input signal *S* (Fig. 1B). Of these, four are tristable and two are bistable. Saddle-node (SN) points in Fig. 1B define transition points between EMT states: SN1, at the boundary of the “low *B*” state, marks the EMT initiation point and SN4, at the boundary of the “high *B*” state, marks the mesenchymal-epithelial transition (MET) initiation point. The tristable states are distinguished by their transition paths and by the accessibility of the intermediate state. Denoting upward transitions (increasing *S*) as “U” and downward transitions (decreasing *S*) as “D”, there can be one or two upward transitions (1U or 2U) and likewise for downward transitions (1D or 2D). By considering both EMT (increasing *S*) and the reverse MET (decreasing *S*), each transition path can be characterized. The 2U2D path passes through the hybrid E/M state during both EMT and MET. The 2U1D path passes through E/M state during EMT but not during MET, i.e. the M state has a larger basin of attraction while that of the E/M state shrinks. Similarly, the 1U2D path passes through E/M state once: during MET but not during EMT. The 1U1D path does not pass through the E/M state in either transition. Finally, there are two bistable EMT landscapes: one is a simple bistable switch with two SN points; the other consists of a double bistable switch. I.e. there are four SN points and a hybrid E/M state exists with a monostable region between two regions of bistability. Of all the EMT landscapes, this has the greatest hybrid state stability, since in this case over some range of *S* only the hybrid state exists.

### Logic controls the effects of direct vs indirect ZEB activation on EMT

To reveal how network logic affects transition paths during EMT and MET, we studied EMT landscapes under perturbations of the direct activation of ZEB (*B*) and the indirect activation of ZEB via inhibition of miR-200 (*A*) for AND vs OR network logic. We perturbed constrained models with symmetrical regulations (i.e. *S* regulates *A* and *B* equally; *A* and *B* inhibit each other equally, etc; for full details see Methods). The direct activation strength (*S_B_*_0_) and the indirect activation strength (*S_A_*_0_) were each perturbed by x% from *S_A_*_0_ = *S_B_*_0_. The resulting EMT landscapes were analyzed for AND (Fig. 2) and OR models (Fig. 3).

**Figure 2:**
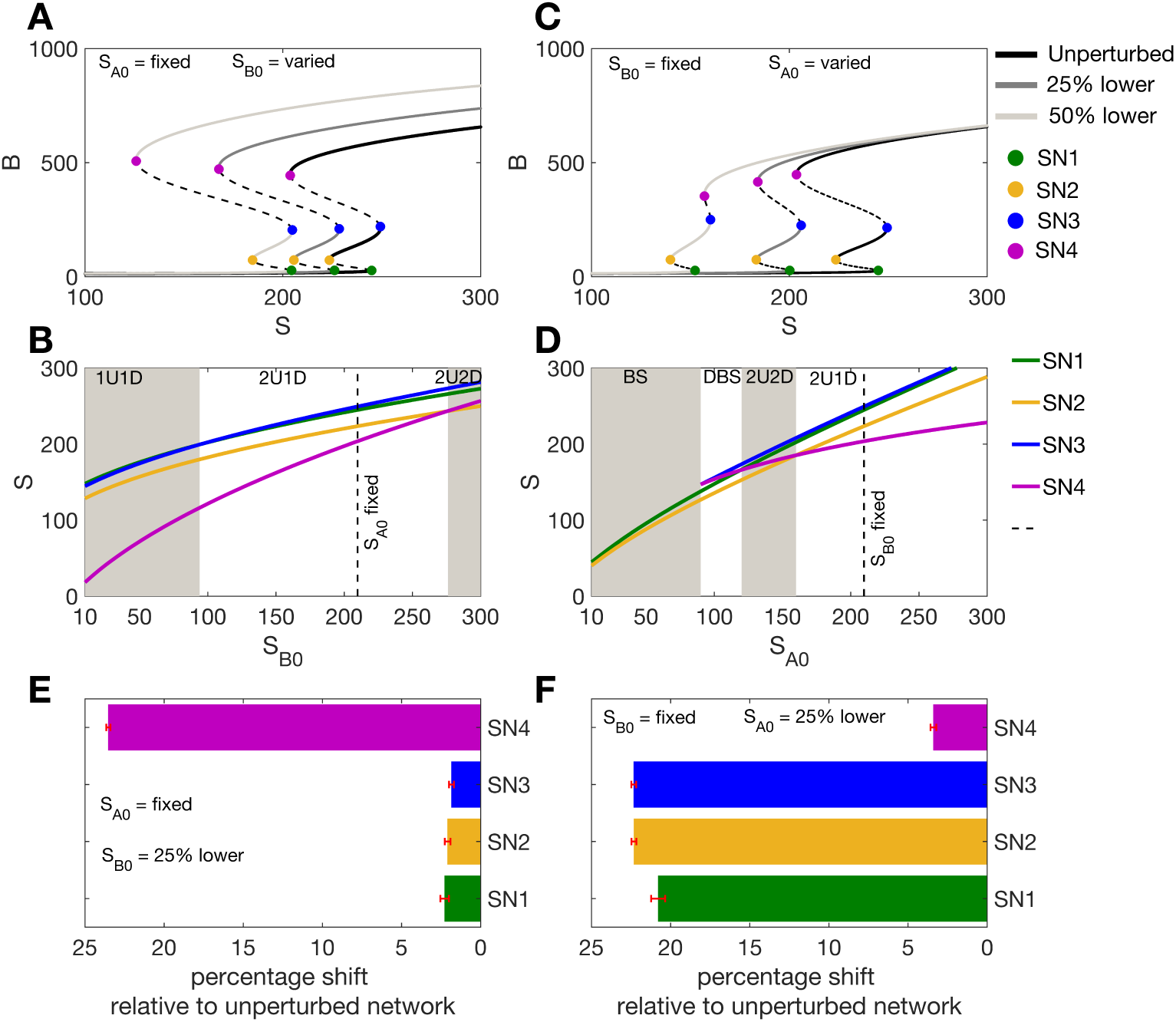
Impacts on EMT of varying regulation strength for AND logic models. **A**. Impact on tristable EMT landscapes for the AND model when varying the direct activation strength (*S*_B0_): unperturbed network and networks perturbed by 25% or 50%. **B**. The loci of the SN points as a function of *S*_B0_. **C**. As for (A), when varying the indirect activation strength (*S*_A0_): unperturbed network and networks perturbed by 25% or 50%. **D**. The loci of the SN points as a function of *S*_A0_. **E**. Sensitivity of the SN points for 144 EMT networks perturbed by a 25% increase in the direct activation strength. **F**. Sensitivity of the SN points for 144 EMT networks perturbed by a 25% increase in the indirect activation strength.

**Figure 3:**
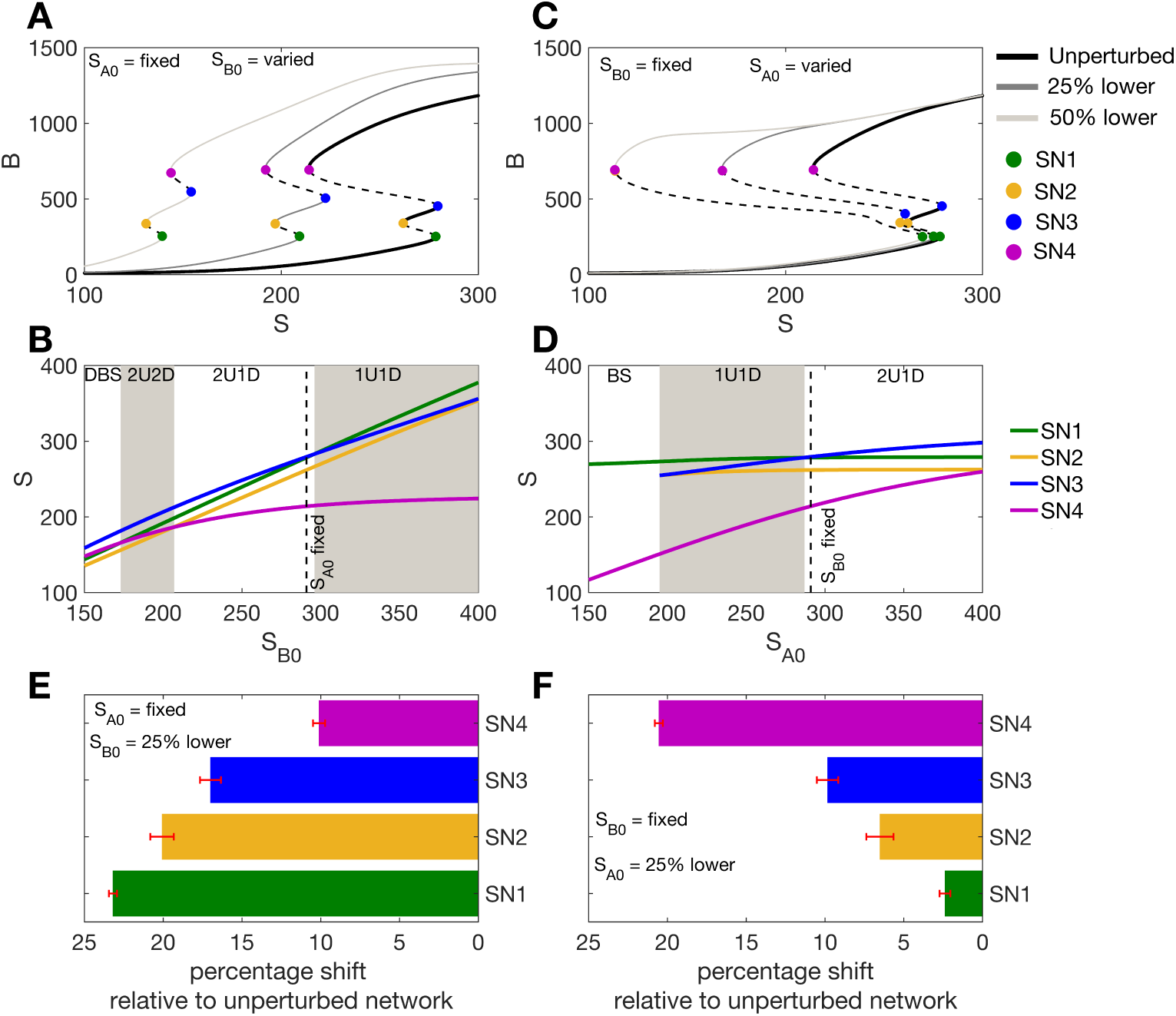
Impacts on EMT of varying regulation strength for OR logic models. **A**. Impact on tristable EMT landscapes for the OR model when varying the direct activation strength (*S*_B0_): unperturbed network and networks perturbed by 25% or 50%. **B**. The loci of the SN points as a function of *S*_B0_. **C**. As for (A), when varying the indirect activation strength (*S*_A0_): unperturbed network and networks perturbed by 25% or 50%. **D**. The loci of the SN points as a function of *S*_A0_. **E**. Sensitivity of the SN points for 108 EMT networks perturbed by a 25% increase in the direct activation strength. **F**. Sensitivity of the SN points for 108 EMT networks perturbed by a 25% increase in the indirect activation strength.

For AND models, we increased the direct (Fig. 2A) or the indirect (Fig. 2C) activation of *B* by 25% from parity (*S_A_*_0_ = *S_B_*_0_; see Methods). Increasing the direct activation rate shifted the whole bifurcation to the left with SN4 (pink) shifting the the most compared to SN1-SN3 (Fig. 2A-B). Fig. 2B depicts the loci of the four SN points as a function of the direct activation strength (*S_B_*_0_). SN4 is most sensitive to changes in the direct activation strength. Increasing the direct activation strength also changes the tristable response type from 2U1D to 1U1D indicating that the hybrid (E/M) state loses stability as the direct activation strength is increased.

When the indirect activation strength *S_A_*_0_ was increased by the same amount, the bifurcation once again shifted to the left, but in this case SN4 changed least. SN1 shifted the farthest and is thus the most sensitive to the indirect activation strength (Fig. 2C-D). Fig. 2D also shows that as the indirect strength increases (lower *S_A_*_0_), the EMT landscape loses tristability and forms a bistable switch. Overall, we see that SN points change according to the perturbation: increasing the indirect activation parameter (Fig. 2C) leads to a much larger change to SN1 than increasing the direct activation parameter by the same amount (Fig. 2A). Thus, EMT is initiated early (with lower *S*) as the indirect regulation strength is increased as compared to an equivalent increase in the direct regulation strength.

The results so far apply for only one set of parameter values. To test their generality, we generated many tristable AND models. To do so we used a method for analysis of model multistability previously described (Dey et al., 2021) (see Methods). For each tristable parameter set, we varied the direct or indirect activation strength by 25% and retained only those parameter sets that generated tristability during both these perturbations. Out of 300 total parameter sets for AND models, 144 permitted tristability under both perturbations. For all 144 tristable parameter sets, we observed that varying the direct activation rate led to large changes in SN4 relative to the other SN points (Fig. 2E), whereas varying the indirect activation rate led to large changes in SN1-SN3 relative to SN4 (Fig. 2F). For the AND model, SN4 is thus most sensitive to changes in the direct activation rate and least sensitive to changes in the indirect activation rate. SN1 on the other hand is most sensitive to changes in the indirect activation rate, suggesting that the indirect activation of ZEB more strongly regulates the initiation of EMT.

An equivalent analysis of direct and indirect activation parameters was performed for the model constructed with OR logic; the results showed opposing effects to those of the AND model. For the OR model, when the direct activation rate was increased by 25% (*S_B_*_0_ lowered), SN1 was the most sensitive and SN4 was the least sensitive point (Fig. 3A-B). When the indirect activation strength was increased by 25% (*S_A_*_0_ lowered), SN4 was the most sensitive, in contrast to the observation for AND models (Fig. 3B,D). Analysis of all paraemter sets permitting tristability after perturbing OR models (108/300 parameter sets) showed that SN1 is most sensitive to the direct activation strength (Fig. 3E) and that SN4 is most sensitive to the indirect activation strength (Fig. 3F). Thus for OR models, the direct activation parameter initiates EMT, since SN1 is most sensitive to this parameter.

Overall, crucial differences emerge regarding the impact of regulatory parameters on EMT depending on the choice of network logic. Changes in SN1 (representing changes in epithelial state stability and the initiation point of EMT) are most sensitive to miR-200 inhibition given AND logic but most sensitive to ZEB activation given OR logic. We see corresponding differences for SN4 (representing mesenchymal state stability and the initiation of MET). Not only is the initiation point of EMT sensitive to different regulations depending on the choice of logic, but different choices of logic can lead one to opposing conclusions about the possible transition paths of cells undergoing EMT.

### Analysis of tristability properties of EMT predicts that the EMT network is wired by AND logic

We studied the properties of tristable models further by analyzing three features of the EMT landscape. The first feature is the saddle node point SN1 (Fig. 1B), SN1 encapsulates the initiation of EMT since EMT is initiated when a transition occurs from the epithelial (E) state into an hybrid (E/M) or mesenchymal (M) state with increasing *S*. Second, for cells in a mesenchymal state, as *S* decreases MET is initiated at SN4. The length of the M state is measured as the distance between SN4 and SN1 (or SN3 depending on the tristable response type). We analyze the M state length here, rather than SN4, as it is more informative than SN4 alone regarding the size of the state space for mesenchymal phenotypes. Third, we analyze the accessibility of the hybrid E/M state, assessed via types of tristable landscape (Fig. 1B). The E/M state is most accessible for 2U2D paths since cells pass through the E/M state during EMT and MET, and least accessible for 1U1D paths since cells do not enter the hybrid state in this case. (This holds in the deterministic case; no longer so when transition paths become probabilistic in the case of stochastic EMT dynamics, which we consider below.)

We analyzed the set of models that permitted tristability after perturbations (144 AND models and 108 OR models) as discussed above. For SN1 (the initiation point of EMT), we saw that increasing the indirect activation strength (i.e. increasing the inhibition of miR200) lowered SN1 (the EMT initiation point) in 96% of the AND models tested (Fig. 4A). I.e. the E state is destabilized and EMT is initiated earlier for increases in the indirect activation relative to the direct activation. For OR models we saw the opposite effect: in 98% of models, SN1 is lowered (EMT initiated earlier) when the *direct* activation strength was increased (Fig. 4B). Thus, network logic dictates the relative importance of regulations on ZEB in initiating EMT. For AND logic, the EMT initiation point is more sensitive to the indirect activation (the inhibition of miR-200 by *S*) than to the direct activation of ZEB. For OR logic, i.e. additive regulation of ZEB, the converse is true: the EMT initiation point is more sensitive to the direct activation of ZEB than to the inhibition of miR-200. In this case, inhibition on miR-200 alone cannot destabilize the E state and initiate EMT without additional direct activation of ZEB.

**Figure 4:**
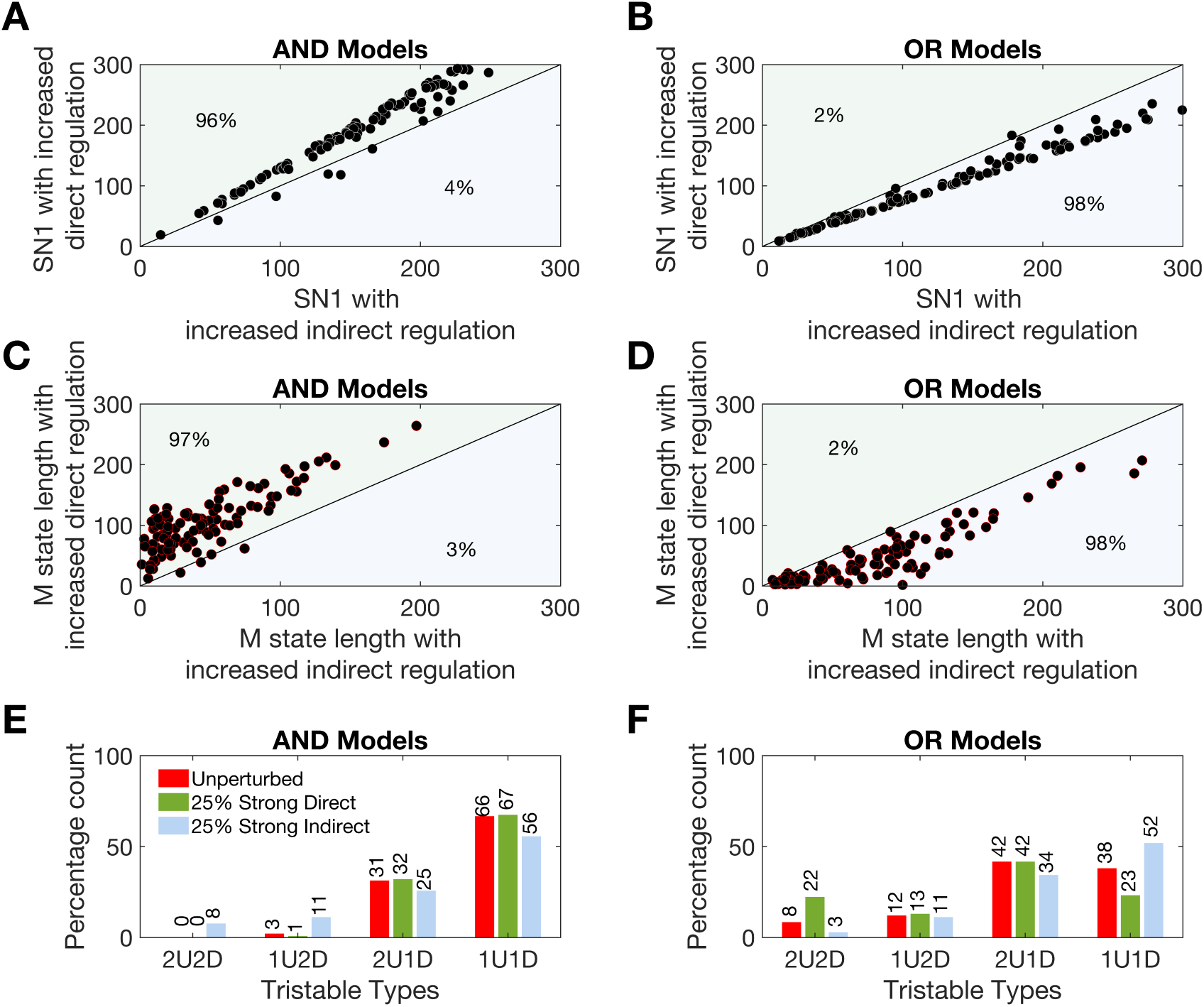
Divergent features of tristable EMT landscapes for AND vs OR models. **A-B**. Scatter plots of SN1, the EMT initiation point, for perturbations to AND models (A) and OR models (B). **C-D**. Scatter plots of the M state length for perturbations to AND models (C) and OR models (D). Solid line drawn for *y* = *x*. **E-F**. The occurrence of different tristable response types for unperturbed models (red bars), direct activation perturbed by 25% (green bars), and indirect activation perturbed by 25% (blue bars) for AND models (E) and OR models (F).

In a similar manner to SN1, we studied how the size of the mesenchymal state attractor (the length of the M state) changed as direct/indirect activation strengths were perturbed for AND and OR models. Here, we found that for AND models increasing the strength of the indirect activation rate decreased the size of the M state (Fig. 4C). I.e. as the inhibition on miR-200 from *S* is reduced, the M state is destabilized and cells in the mesenchymal state more readily undergo MET. This is supported by experimental evidence: miR-200 is known to maintain the E state such that suppressing is sufficient to initiate EMT, and re-expression it is sufficient to induce MET (Korpal et al., 2008; Nagai et al., 2024). Analysis of OR models showed the opposite behaviour: increasing the indirect activation (reducing the suppression on miR-200) increased the length of the M state (Fig. 4D), i.e. increasing the likely population size of mesenchymal cells.

We also analyzed how accessibility of the hybrid state was affected by perturbing the direct and indirect regulation rates. For AND models, the accessibility of the hybrid state increased (more 2U2D paths and fewer 1U1D paths) as the indirect activation rate increased (Fig. 4E). For OR models — as seen above — the converse is true: the accessibility of the hybrid state increased as the direct activation rate increased (Fig. 4F). The observation for AND models is supported by experimental studies where, in MDA-MB-231 cells, it was shown that the induction of miR-200 promoted cells to enter into a hybrid E/M state displaying a hybrid phenotype characterized by collective cell migration and epithelial gene expression (Nagai et al., 2024).

Similar results were obtained when the direct/indirect regulation parameters were perturbed by larger amounts: 50% from the baseline (Fig. S1). We also tested an alternate indirect model parameter on EMT phenotypes. Two interactions in the GRN combine to define the “indirect activation of ZEB:” the inhibition from *S* to *A* (*S_A_*_0_; explored above), and the interaction from *A* to *B* (*A_B_*_0_). Perturbation analysis performed for *A_B_*_0_ generated similar qualitative results for AND and OR models as seen above: where the logic dictates which network perturbation will control specific EMT phenotypes (Fig. S2).

In summary, through perturbation of direct/indirect regulation parameters, alternate EMT states are stabilized or destabilized based on the choice of network logic. For AND models, increasing the inhibition on miR-200 (thus increasing the indirect activation of ZEB) destabilized the accessibility of the epithelial &mesenchymal states and increased the accessibility of the hybrid state. In contrast, for OR models, increasing the inhibition on miR-200 stabilized the epithelial &mesenchymal states and delayed the initiation of EMT. The hybrid E/M state was in this case less accessible. Overall, we saw that when the E state is destabilized, so does the M state and the accessibility of the hybrid state increases, and vice versa. A considerable body of literature on the role of miR-200 in the initiation of EMT and the reverse MET suggests that suppressing miR-200 is sufficient to initiate EMT, and that re-expressing miR-200 in mesenchymal state cells can initiate MET (Kong et al., 2009; Korpal et al., 2008; Nagai et al., 2024; Zhang, Liu, et al., 2012). In light of these experimental studies and our results on the alternative phenotypes observed with network logic, we predict that the GRN characterizing EMT via regulation of ZEB by miR-200 and SNAIL is constructed with AND logic.

### The role of network logic is conserved for a wide family of models

Until this point, we have perturbed parameters by fixed amounts from a baseline model. This allowed us to quantify how EMT is differentially regulated in comparable scenarios, as the baseline model was constrained to have equal production/degradation rates and equal interaction strengths between nodes. However, it is also important to assess whether results obtained hold true more broadly. To test this, we relax the constraints previously imposed, and consider a wide range of parameterized EMT network models that permit tristability (Table 1; “unconstrained models”). For models constructed with either AND or OR logic, we sampled parameters and analyzed properties of the EMT landscape. Through systematic perturbations of the direct and indirect regulations on ZEB for unconstrained models, we observed the same qualitative results as obtained above for constrained models. The EMT initiation point, stability of the mesenchymal state, and accessibility of the hybrid state all exhibit sensitivity to the direct/indirect regulation parameters that is dictated by the network logic (Fig. S3).

**Table 1:**
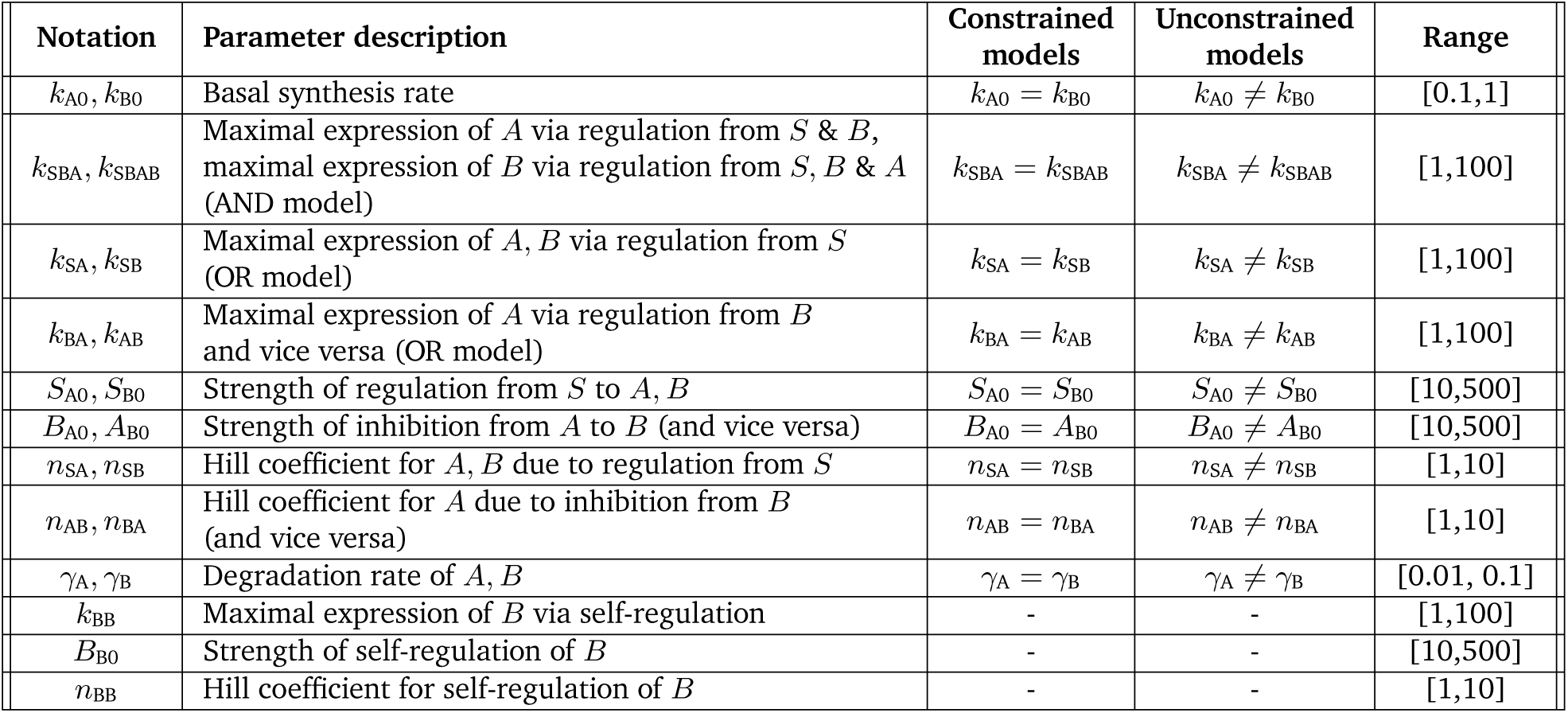
Description of model parameters and values. The first column gives the parameter notation, the second column is the parameter description, the third and fourth columns give relationships between parameters under different models, and the fifth column gives the parameter range used for sampling. Hill coefficients take integer values; all other parameters take real values.

We also analyzed the parameters of a different gene regulatory network model of EMT. The ternary chimera switch model (Lu et al., 2013) is constructed with AND logic and has a similar form to Eqns. 1 but with a larger number of parameters. For the Lu et al. model, we analyzed the bifurcation landscapes for ZEB mRNA and ZEB protein with respect to SNAIL as the direct or indirect activation strengths were increased by 55.5% (Fig. S4). In agreement with the analysis of AND models above, increasing the indirect activation strength in the Lu et al. model lowered the initiation point for EMT, increased the accessibility of the E/M state, and destabilized the M state (Fig. S4). We sought to perform a similar analysis for the model of Tian et al. (2013) that is constructed with OR logic and characterized by two cascading bistable switches. However, this model is constructed differently in that different modules of the network control different parts of the cascading bistable switches. There is therefore no obvious way in which direct vs. indirect regulations of an EMT initiating factor can be defined.

We also investigated the impact of other model parameters on the EMT GRN. Perturbation analysis for the self-activation of *B* (*B_B_*_0_) and the inhibition on *A* from *B* (*B_A_*_0_) was performed for constrained models, revealing that increasing the self-activation of *B* (decreasing *B_B_*_0_) increased the length of the E/M state in all 144 tristable AND models (Fig. S5A). This observation is in agreement with previous work (Lu et al., 2013), and indeed is a general phenomenon observed for the self-activation rate in tristable networks of this type (Jia, Jolly, Harrison, et al., 2017). For OR models, in the majority (67%) of cases we saw that increasing the self-activation of *B* (decreasing *B_B_*_0_) increased the stability of the E/M state (Fig. S5B). For both AND and OR models, increasing self-activation (decreasing *B_B_*_0_) increased the accessibility of the hybrid E/M state (Fig. S5C-D). SN1 also behaved similarly with respect to *B_B_*_0_ for AND and OR models (Fig. S5E-F). For the inhibition on *A* from *B* (*B_A_*_0_): decreasing the inhibition strength led to increased accessibility/stability of the hybrid state for both AND and OR models (Fig. S6). This demonstrates that — unlike for the direct/indirect activation rates of ZEB — perturbing the self-activation rate of *B* or the inhibition on *A* from *B* leads to consistent phenotypes regardless of the network logic with which the model is constructed.

### Network logic and the combinatorics of gene regulation

In light of the opposing effects of network logic on EMT phenotypes observed for a wide variety of tristable models, we explored how different responses to the same parameter perturbations could be explained in terms of combinatorial gene regulation. The node *B* (ZEB in the EMT model) receives in total three inputs: self-activation, activation from *S* (direct activation), and inhibition from *A* (indirect activation by *S*). For AND models, input regulations to *B* combine multiplicatively. This implies a strong dependence: the direct activation of *B* only weakly affects EMT initiation when there remains inhibition on *B* from *A*. Thus for AND models, *B* is less sensitive to direct activation from *S*. In constrast, increasing the indirect activation rate weakens the inhibition from *A* on *B*. This implies a lower activation threshold of *B*, making it more sensitive to changes in *S* (earlier initiation of EMT). Increasing the indirect activation strength also destabilizes the M state, thus decreasing the EMT window: the length between SN1 and SN4, i.e. total size of the multistability region (Fig. S7A).

For OR models, input regulations to *B* combine additively, such that each individually regulates *B*. This relative independence means that even if one of the regulations is low/off, *B* can still be expressed. In this case, increasing the direct activation rate has a strong effect on *B*, since *S* can activate *B* even in the presence of inhibition from *A*. Earlier EMT initiation (lower SN1) reduces the distance between SN1 and SN4, thus reducing the EMT window. For the OR model, increasing the indirect activation rate (less inhibition from *A* to *B*) has a weak effect on *B*, since an increase in *S* (direct activation) is still required to initiate EMT. Since in this case *B* is less sensitive to *S*, also for the reverse MET a greater reduction in *S* is required to initiate the transition. Thus increasing the indirect activation rate increases the EMT window (Fig. S7B). By teasing apart the combinatorial effects of additive vs. multiplicative regulation on GRN interactions, the relative importance of specific model parameters can thus be deduced. In a larger EMT network that includes mRNA and protein species of ZEB (as in Tian et al. (2013)), it is no longer possible to clearly define a direct activation. However, considering the two possible paths, we see that the path more closely related to direct activation (via ZEB mRNA) behaves similarly to previous results for AND vs OR models, albeit less distinctly: AND and OR phenotypes show more overlap (Fig. S8).

### Experimental design can reveal the network logic in cells undergoing EMT

The opposing effects of AND vs OR logic on EMT phenotypes can be analyzed in light of data to inform experimental design and reveal how EMT transcriptional networks are wired in specific cell types. Zhang, Donaher, et al. (2022) used a transformed human mammary epithelial cell line (HMLER) to create single-cell clones that displayed markedly varied EMT phenotypes when maintained in culture. Approximately 3/4 of the single-cell-derived clones were non-convertible: they stably maintained their epithelial status in culture. In contrast, 1/4 of the clones were convertible: they were able to spontaneously undergo EMT and displayed a spectrum of EMT states including hybrid E/M cells.

To compare model-predicted EMT phenotypes with the tristable responses observed in live HMLER cells (Zhang, Donaher, et al., 2022), we studied stochastic EMT transition paths for the miR200—ZEB network simulated via stochastic differential equations (see Methods). SDEs were specified by an additive noise model to constrain the EMT cell fate choices to be consistent with those for the deterministic model, since more complicated noise models can alter the cell fates available on a phenotypic landscape (Coomer et al., 2021). For low levels of signal *S*, cells cannot transition out of an E state, i.e. these cells are non-convertible (Fig. 5A [red box]). For sufficiently high levels of *S*, cells are able to undergo EMT and produce a range of EMT states, i.e. corresponding to convertible clones (Fig. 5A [blue box]). From analysis of the EMT transition landscapes at different levels of input stimulus (*S_i_*) we can obtain the point at which each population is able to transition out of the E state and undergo EMT. For the unperturbed network, cells first transition at *S_u_* = *S*_6_, and the mesenchymal state becomes accessible at *S*_7_ (Fig. 5B).

**Figure 5:**
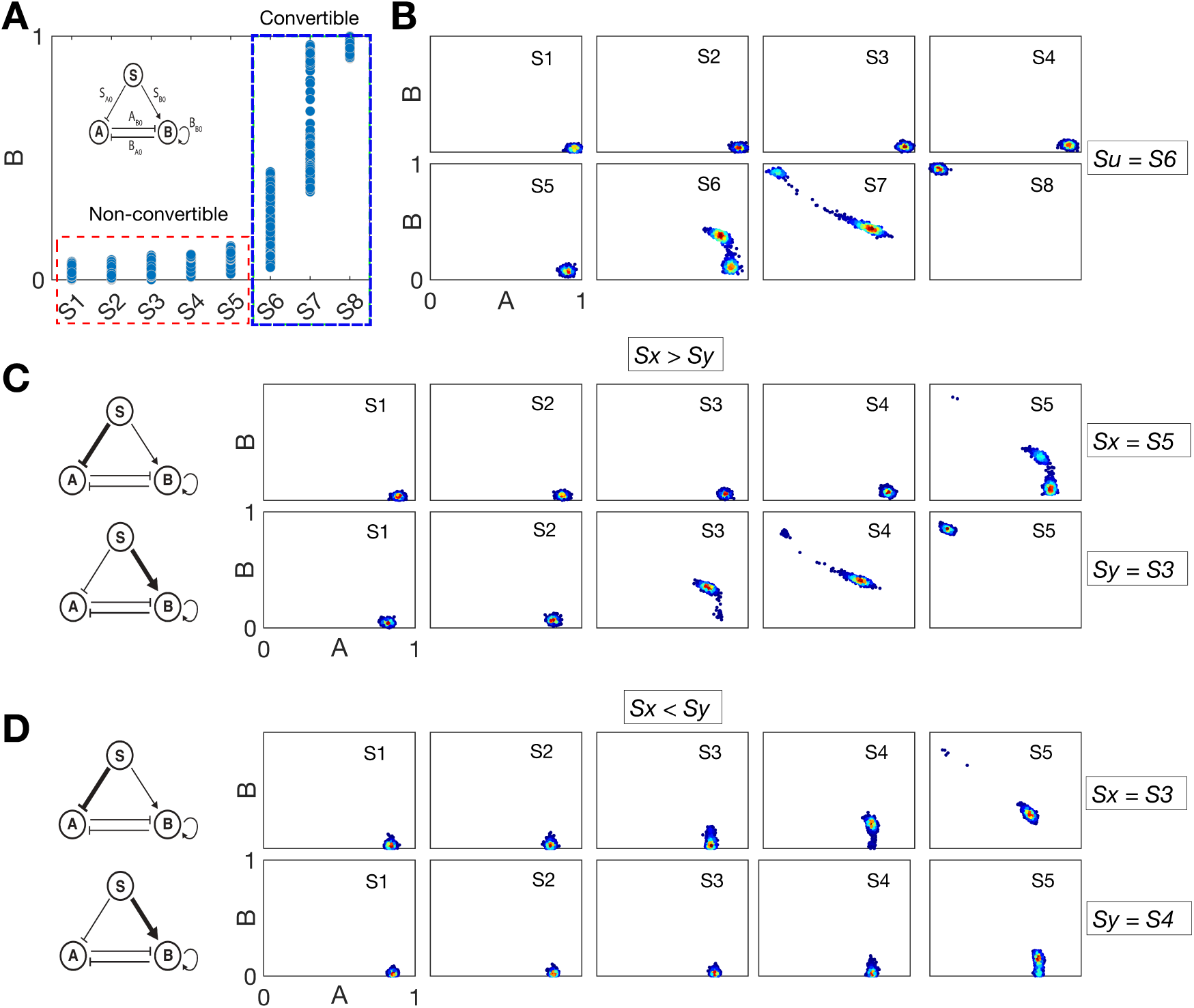
Experimental design reveals the network logic of GRNs for EMT. **A**. Stochastic simulations of the EMT GRN network for different *S* values can categorize cells as non-convertible (signal levels *S* = *S*_1_–*S*_5_; red box) or convertible (signal levels *S* ≥ *S*_6_; blue box). **B.** Equilibria of the model in the (*A, B*) phase plane for stochastic differential equation simulations of the unperturbed model. *S_i_* indicates an input signal strength of signal *S*. **C.** Equilibria of the model in the (*A, B*) phase plane for simulations of the OR logic GRN. Either the indirect (top row) or the direct (bottom row) regulation on ZEB is perturbed and the minimum signal required to initiate EMT is recorded (*S_x_*for indirect and *S_y_*for direct). **D.** Equilibria of the model in the (*A, B*) phase plane for simulations of the AND logic GRN, perturbing either the indirect or the direct regulation as for (C).

Stochastic simulation of EMT networks constructed with AND vs. OR logic show markedly different transition paths, allowing us to design experiments that can infer the EMT network logic that is present in HMLER cells. From a baseline transition point obtained by simulating the unperturbed model (*S_u_* = *S*_6_; Fig. 5B), we study how the EMT transition landscape changes as the direct or the indirect ZEB activation rates are perturbed. The dose of stimulus at which cells first transition when the inhibition of miR-200 is increased (increased indirect activation) is recorded as *S_x_*. The dose of stimulus at which cells first transition when the direct activation of ZEB is increased is recorded as *S_y_*. For models constructed with OR logic, we see that *S_x_* = *S*_5_ and *S_y_* = *S*_3_, i.e. *S_x_ > S_y_* (Fig. 5C). For models constructed with AND logic, we see that *S_x_* = *S*_3_ and *S_y_*= *S*_4_, i.e. *S_x_ < S_y_* (Fig. 5D). These inequalities offer means with which to determine the network logic in live cells: if *S_x_ > S_y_*, then the network is constructed with OR logic; if *S_x_ < S_y_*, then the network is constructed with AND logic.

Sufficient experimental data to discern between these scenarios can be gathered by subjecting non-convertible HMLER clones to different levels of stimulus via SNAIL or TGF-*β* under two separate perturbations. The first would be the inhibition of miR-200: this could be achieved by steric blocking; the second would be an amplification of the rate of ZEB activation via SNAIL (which could be achieved by epigenetic regulation or the addition of SNAIL/ZEB interacting co-factors, such as EGR1 or SP1 (Wu et al., 2017)). By virtue of the inequalities derived above, neither knowledge of the precise model parameterization nor of the precise stimulus levels matching simulations are required for inference. By determining the relative points at which cells become convertible, the network logic can be inferred. Observing EMT earlier when miR-200 is inhibited than when ZEB activation is increased implies that the network is constructed with AND logic, where signals combine multiplicatively. Observing the converse implies that the network is constructed with OR logic, where signals combine additively.

## Discussion

We have developed a GRN model of EMT to study the effects of network logic on cell state transition paths. The GRN describes the regulation of ZEB by two distinct routes: direct activation of ZEB by SNAIL or indirect activation of ZEB through a double inhibition: SNAIL −1 miR-200 −1 ZEB (Barrallo-Gimeno et al., 2005; Kaufhold et al., 2014; Sánchez-Tilló et al., 2010; Zhang, Sun, et al., 2015). The presence of two coupled positive feedback loops leads to tristable landscapes for this GRN. Through bifurcation and statistical analyses, we have shown that the choice of network logic (multiplicative vs. additive) is important: different choices of logic lead to divergent EMT phenotypes. We have also demonstrated how model predictions can be used to design experiments to discriminate between EMT network, which could thus be used to infer the network logic of the network in live cells.

Current literature characterizing the role of miR-200 in EMT has shown that when miR-200 is inhibited, ZEB transcription factors are upregulated (Díaz-López et al., 2014; Gregory et al., 2008; Hill et al., 2013; Klicka et al., 2022; Korpal et al., 2008; Title et al., 2018). Moreover, re-expression of miR-200 can cause a loss of mesenchymal traits and a reverse MET transition to an epithelial state (Kong et al., 2009; Korpal et al., 2008; Nagai et al., 2024; Zhang, Liu, et al., 2012). These data are most consistent with EMT network models constructed with multiplicative logic, i.e. where SNAIL and miR-200 act cooperatively to regulate EMT. In agreement with this prediction, recent studies in a mesenchymal cell line have found that the induction of miR-200 cells causes cells to transition into a hybrid E/M state that promotes collective cell migration and expresses epithelial genes (Nagai et al., 2024), although it may be possible to design strategies to avoid hybrid states during MET (Kim et al., 2023).

Divergent EMT phenotypes were observed by perturbing two regulation parameters: the direct activation of ZEB by SNAIL and the inhibition of miR-200 by SNAIL. Unlike the self-activation of ZEB, these two parameters were highly sensitive to the combinatorics of gene regulation. The relevance of these results extends beyond EMT: the network structure of the GRN is found in a variety of biological systems (Frankhouser et al., 2024; Mangan and Alon, 2003). It is notable that experimental data are unlikely to help with the general challenge of network logic choice in modeling, since genes exhibit a wide range of responses to multiple inputs, from additive to multiplicative, as well as sub-/super-additive and multiplicative responses (Sanford et al., 2020).

There are limitations to the current work to be addressed in future studies. The three-node EMT network studied (a coherent feedforward loop between SNAIL, miR-200, and ZEB) does not consider the transcriptional species of SNAIL or ZEB, unlike alternative EMT models (Tian et al., 2013). Transcriptional and translational regulation are considered jointly here for parsimonious modeling. If mRNA species were modeled, the state space would expand to five dimensions such that would no longer be a simple feedforward loop and thus it would be unclear how to discern direct vs indirect regulation of ZEB (Fig. S8). Nonetheless, future extensions of the work ought to consider these more sophisticated regulatory paths. While a substantial body of literature supports the interactions in this EMT network (Díaz-López et al., 2014; Gregory et al., 2008; Hill et al., 2013; Klicka et al., 2022; Korpal et al., 2008; Tian et al., 2013; Title et al., 2018; Zhang, Tian, et al., 2014), there are certainly additional factors that regulate EMT and could change the dynamical landscape. These include epigenetic factors (Al-Radhawi et al., 2022) as well as larger EMT networks that permit tetrastability (Hong, Watanabe, et al., 2015; Kim et al., 2023). In such a case two hybrid E/M states exist, giving rise six saddle points and subsequently many more possible transition paths. Additional hybrid states have clinical implications: if on such an EMT landscape cells can more readily access one or more hybrid states, this could increase metastatic potential. The effects that network logic mediate with larger EMT networks or in light of epigenetic regulatory factors on multistable landscapes will be an important direction for future work.

The structure of the network (consisting of species *S*, *A*, and *B* in its general form) defines a coherent feedforward loop; such network motifs are found in a variety of biological systems (Kalir et al., 2005; Mangan, Zaslaver, et al., 2003; Pieters et al., 2021). The central result of this work that network logic dictates the relative impact of network regulations on phenotype applies in the context of this particular network motif. It would be interesting to extend the analysis of network logic to other network motifs of three or four interacting species (Alon, 2020; Mayer et al., 2023). However, as discussed above, in larger networks it becomes more challenging to distinguish direct vs. indirect regulations. The changes in phenotype observed required perturbation of at least two network edges, i.e. a systems biology approach is inherently required (single perturbations are insufficient). This complicates experimentation, although as we outline it will still be possible to distinguish effects via experiments, e.g. in HMLER cells (Zhang, Donaher, et al., 2022) by first perturbing miR-200 inhibition and then activation of ZEB. Analysis of EMT transition paths required inference of the landscapes from the GRN. Alternative landscape-based approaches do not require knowledge of the specific GRN and offer means to study transition paths when the GRN is not known (Boukacem et al., 2024; Liu et al., 2023; Seyboldt et al., 2022).

Through analysis of how specific regulations in a GRN control phenotypes, we have explained how network logic dictates phenotypes, and offer means with which to infer the EMT network logic in cells through experimentation. In the case of EMT in cancer metastasis, this will help us to design interventions that decrease the probability that cells undergo EMT or form hybrid E/M cell states with metastatic potential. Through analysis of general network properties, we have shown how small logical gene circuits can be constructed to achieve desired outcomes in cells that experience different perturbations, of relevance for synthetic biology and the design of gene networks *in silico*. The ability to explain how multiple signals regulate their target genes offers new means with which to understand the dynamics of gene regulatory networks and the cells fate decisions they control.

## Materials and Methods

### A three-node GRN model of EMT

We study a three-node GRN model that characterizes EMT (Fig. 1A). This network consists of an input signal SNAIL (*S*) that regulates miR-200 (*A*) and ZEB (*B*). We investigate two mathematical models of this network that differ in their network logic.

The model derived from this EMT network constructed with AND logic is given by the equations:

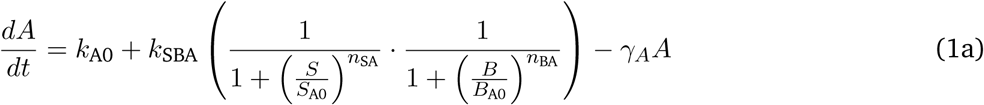

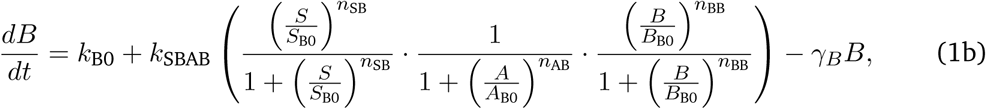

where the parameter descriptions and values used are given in Table 1.

The model derived from this EMT network constructed with OR logic is given by the equations:

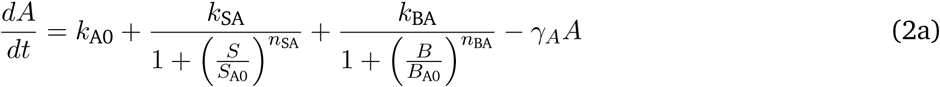

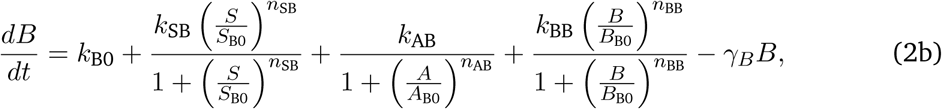

where the parameter descriptions and values used are given in Table 1. The parameters *k*_A0_*, k*_B0_ are associated with the basal synthesis rate of *A* and *B*, respectively. The maximal expression rate of *A* due to *S* and *B* is represented by *k*_SBA_ for AND model and by *k*_SA_ and *k*_BA_, respectively for the OR model. The parameters *S*_A0_*, B*_A0_ are associated with threshold values of *S* and *B* at half-maximal value of *A*, respectively. The Hill coefficients on *A* due to *S* and *B* are given by *n*_SA_ and *n*_BA_, respectively. The maximal expression rate of *B* due to *S*, *A* and *B* is represented by *k*_SBAB_ for AND model and by *k*_SB_*, k*_AB_ and *k*_BB_, respectively for OR model. The parameters *S*_B0_*, A*_B0_ and *B*_B0_ are associated with threshold values of *S*, *A* and *B* at half-maximal value of *B*, respectively. The Hill coefficients for regulation of *B* via *S*, *A* and *B* are *n*_SB_*, n*_AB_ and *n*_BB_. The parameters *γ_A_, γ_B_* represent the degradation of *A* and *B*.

### Generation of tristable EMT models

To generate many tristable models with either AND or OR logic, we followed the automated bifurcation generator method developed by Dey et al. (2021), in which the multi-dimensional ODE system is reduced to a one dimensional ODE using a transfer function.

Briefly, given the ODEs for the AND model:

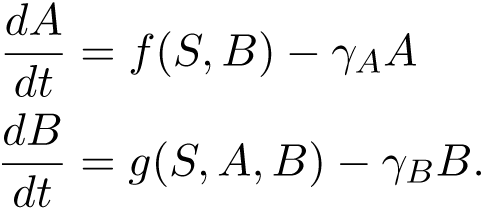

At steady state, 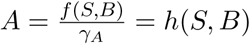 and we have

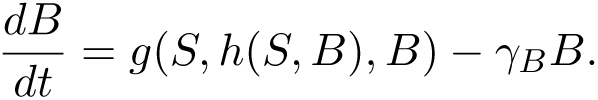

The effective force *F* (*S, B*) of the system is thus:

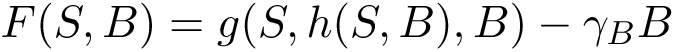

And the effective potential *V* (*S, B*) is given by:

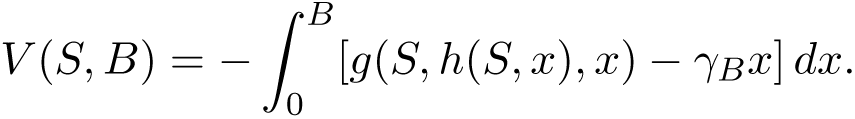

The effective potential is then used to plot potential landscapes for each model. Potential landscapes are plotted in the range *B* ∈ [0, 3000] and *S* ∈ [0, 300]; all other parameters were sampled from uniform distributions over the ranges specified in Table 1. For a sampled parameter set, potential landscapes are generated for many values of *S* ∼ Unif(0, 300), and for each landscape the total number of local minima/maxima are recorded. If at any value of *S* the potential energy landscape contains 3 local minima and 2 local maxima, the parameter set is recorded as permitting tristability. Sampling continued until a total of 300 parameter sets permitting tristability were recorded for each of the AND and the OR models.

### Perturbation analysis of tristable EMT models

Comparisons of phenotypes on the EMT landscapes were performed by perturbing tristable EMT models in the following manner. Given a parameter set that permits tristability, and the corresponding bifurcation plot (the unperturbed EMT landscape *U*), we investigate the effects of perturbing two parameters: the inhibition by *S* on miR-200 (indirect activation parameter: *S*_A0_) and the direct activation of ZEB by *S* (the direct activation parameter: *S*_B0_). I.e.

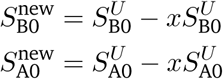

where *S^U^*denotes the unperturbed parameter and we consider perturbations of size *x* = 0.25 and *x* = 0.5. Note that, due to the form of the model, we refer to a *decrease* in the parameter value by *x*% as an *increase* in the strength of regulation via that parameter by *x*%. All other parameters remain unchanged, ans we compare the unperturbed landscapes with the landscapes generated by 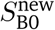 or 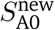.

Perturbation analysis is performed for both the constrained and the unconstrained models (defined in Table 1). For the constrained models, 300 tristable models as described above were perturbed and their potential energy landscapes were generated. 144 AND models and 108 OR models permitted tristability following perturbation of both *S*_B0_ and *S*_A0_ by 25%.

### Analysis of EMT landscape properties: SN sensitivity, M state attractor size, and bifurcation type

We assess the sensitivity of a saddle node (SN) point to a change in parameter using the mean shift in value of that SN point for a set of landscapes for perturbed models relative to their unperturbed counterparts. via:

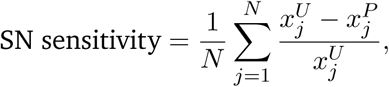

where *x^U^*are the SN point values for unperturbed models and *x^P^* are the SN point values for the perturbed models. *j* = 144 models for AND logic; *j* = 108 models for OR logic. Each of the four SN points is evaluated in the same way.

The size of attractors for the mesenchymal (M) state is quantified by the length of the M state. The length of the M state is measured for *S* as the distance between SN4 and max(SN1,SN3).

The tristable bifurcation type is categorized according to the criteria in Fig. 1B, i.e. for tristable models there are four possible bifurcation types. Ranked from the type permitting the fewest hybrid states to the type permitting the most, these are: 1U1D, 1U2D = 2U1D, 2U2D. For the total number of tristable networks in a given set, the fraction of each bifurcation type is recorded for the EMT landscape under different perturbations of the network.

### Stochastic simulations of EMT GRN models

Stochastic simulations of EMT network models are performed using a stochastic differential equation (SDE) formulation of each model. I.e., given a general ODE model as 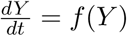 for a vector of species *Y*, we express the stochastic GRN dynamics in the form:

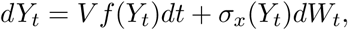

where *f* denotes the EMT network interactions and takes the same form as for the ODE equivalent, *σ* represents the noise model, and *dW_t_* denotes an increment of the Weiner process. *V* is a volumetric scaling factor. The SDE model can be solved numerically using the Euler-Maruyama method, where we assume an Itô interpretation.

The scaling factor *V* is introduced to scale the values of species *A* and *B* into a similar range. This is done such that for models with equal magnitude of the noise (*σ_x_* = 1), stochastic fluctuations will occur on similar scales of magnitude.

SDE models constructed with different logic, i.e. with *f* (*Y_t_*) specified by the RHS of either Eqs. 1 (AND logic) or Eqs. 2 (OR logic), are simulated for 1000 time steps with a step size of 0.01, which we found appropriate to approximate the stationary state. The final values *Y*_end_ = (*A*^∗^*, B*^∗^) are considered to be the steady state values. To simulate a population of single cells, we assume that each simulation represents a cell, and consider a population of *N* = 1000 cells, i.e. for every value of *S* considered, 1000 trajectories are simulated to obtain a final population of cells 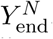. Stochastic simulations are performed at eight values of *S*, i.e. there are a total of 8*N* cells. Steady state values of *A* and *B* are min-max normalized. I.e. within the total set of 8N simulations:

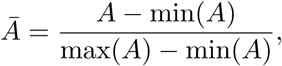

with *B̅* calculated similarly.

## Acknowledgements

This work was supported by the National Institutes of Health (NIH) R35GM143019 to A.L.M.

## Author Contributions

**A.D.**: Conceptualization, software, methodology, investigation, formal analysis, writing— original draft, writing—reviewing & editing. **A.L.M.**: Conceptualization, methodology, investigation, formal analysis, supervision, writing—original draft, writing—reviewing & editing.

## Data Availability

All of the code and analysis scripts associated with this manuscript were developed in MATLAB (Natick, MA) and are available on GitHub at: https://github.com/maclean-lab/network-logic-EMT.

