## Supplementary Figures for "Transition paths across the EMT landscape are dictated by network logic"

### Supplementary Figures for
### Transition paths across the EMT landscape are dictated by transcriptional network logic

Anupam Dey[1], Adam L. MacLean[1,*]
[1]Department of Quantitative and Computational Biology, University of Southern California, Los Angeles, CA

#### Contents

**Supplementary Figure 1** **3**

**Supplementary Figure 2** **4**

**Supplementary Figure 3** **5**

**Supplementary Figure 4** **6**

**Supplementary Figure 5** **7**

**Supplementary Figure 6** **8**

**Supplementary Figure 7** **9**

**Supplementary Figure 8** **9**

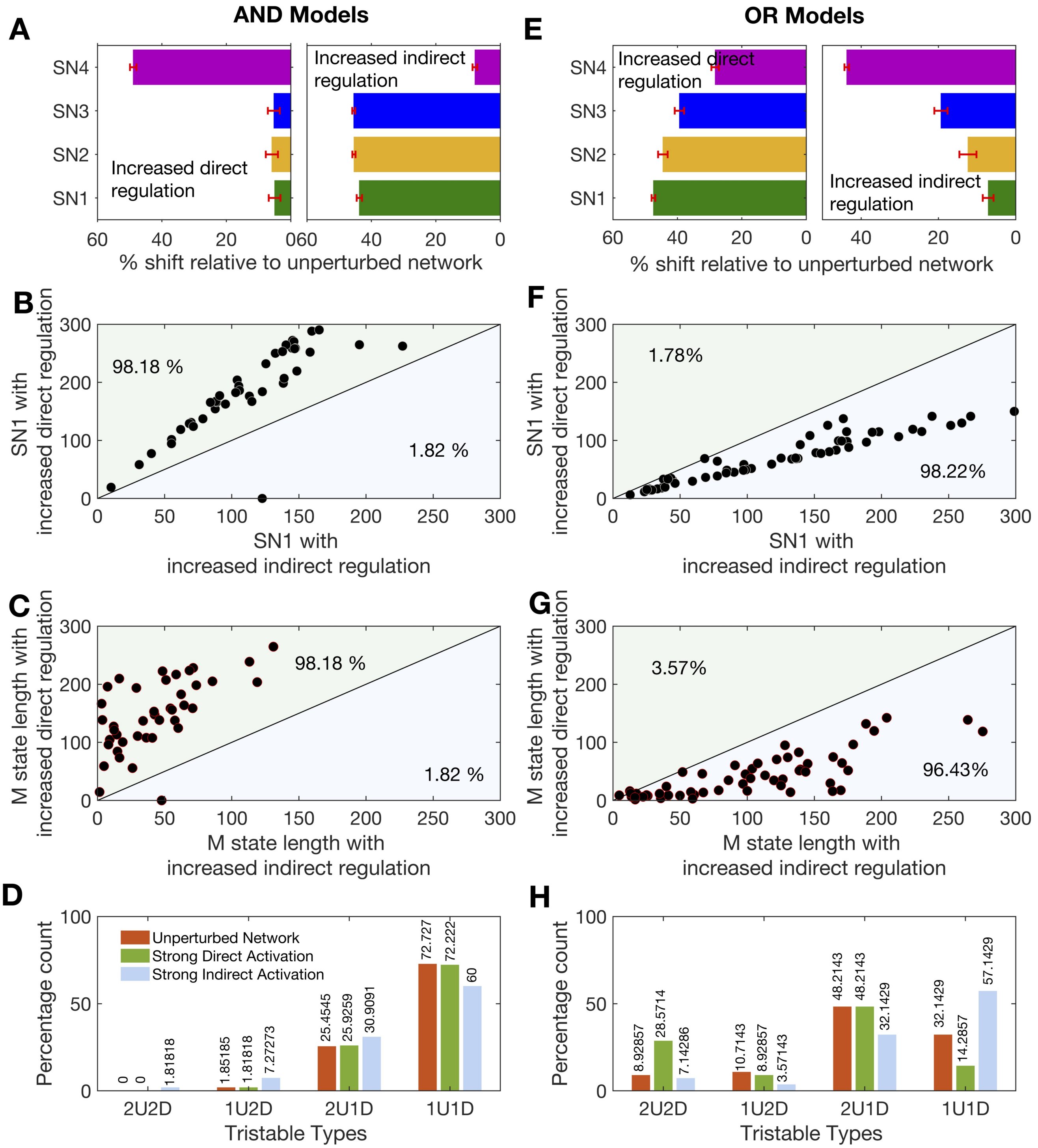


**S1 Figure.** Impact of varying regulation parameters by 50% in AND/OR models. **A**. Sensitivity of the SN points for tristable AND models perturbed by a 50% increase in the direct or the indirect activation strength. **B**. Scatter plots of SN1 (the EMT initiation point) for 50% perturbations to AND models. **C**. Scatter plots of the M state length for 50% perturbations to AND models. **D**. The occurrence of different tristable response types for unperturbed models (red bars), direct activation perturbed by 50% (green bars), and indirect activation perturbed by 50% (blue bars) for AND models. **E**. Sensitivity of the SN points for tristable OR models perturbed by a 50% increase in the direct or the indirect activation strength. **F**. Scatter plots of SN1 (the EMT initiation point) for 50% perturbations to OR models. **G**. Scatter plots of the M state length for 50% perturbations to OR models. **H**. The occurrence of different tristable response types for unperturbed models (red bars), direct activation perturbed by 50% (green bars), and indirect activation perturbed by 50% (blue bars) for OR models.

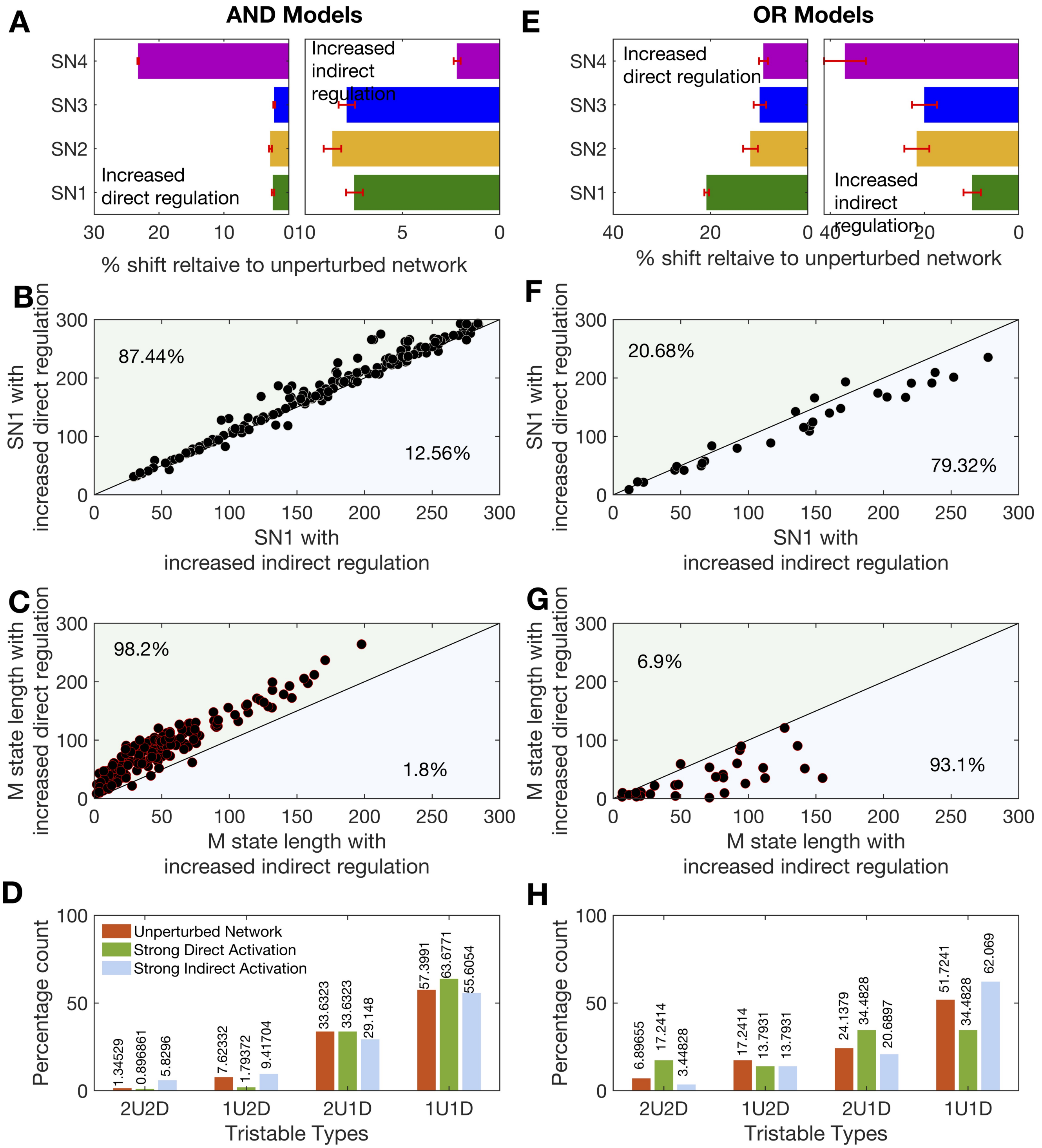


**S2 Figure.** Impact of varying alternative indirect regulation parameter $A_{B0}$ in AND/OR models. **A**. Sensitivity of the SN points for tristable AND models perturbed by a 25% increase in the direct activation strength or the *alternative indirect activation parameter* $A_{B0}$. **B**. Scatter plots of SN1 (the EMT initiation point) for 25% perturbations to AND models. **C**. Scatter plots of the M state length for 25% perturbations to AND models. **D**. The occurrence of different tristable response types for unperturbed models (red bars), direct activation perturbed by 25% (green bars), and indirect activation perturbed by 25% (blue bars) for AND models. **E**. Sensitivity of the SN points for tristable OR models perturbed by a 25% increase in the direct activation strength or the *alternative indirect activation parameter* $A_{B0}$. **F**. Scatter plots of SN1 (the EMT initiation point) for 25% perturbations to OR models. **G**. Scatter plots of the M state length for 25% perturbations to OR models. **H**. The occurrence of different tristable response types for unperturbed models (red bars), direct activation perturbed by 25% (green bars), and indirect activation perturbed by 25% (blue bars) for OR models.

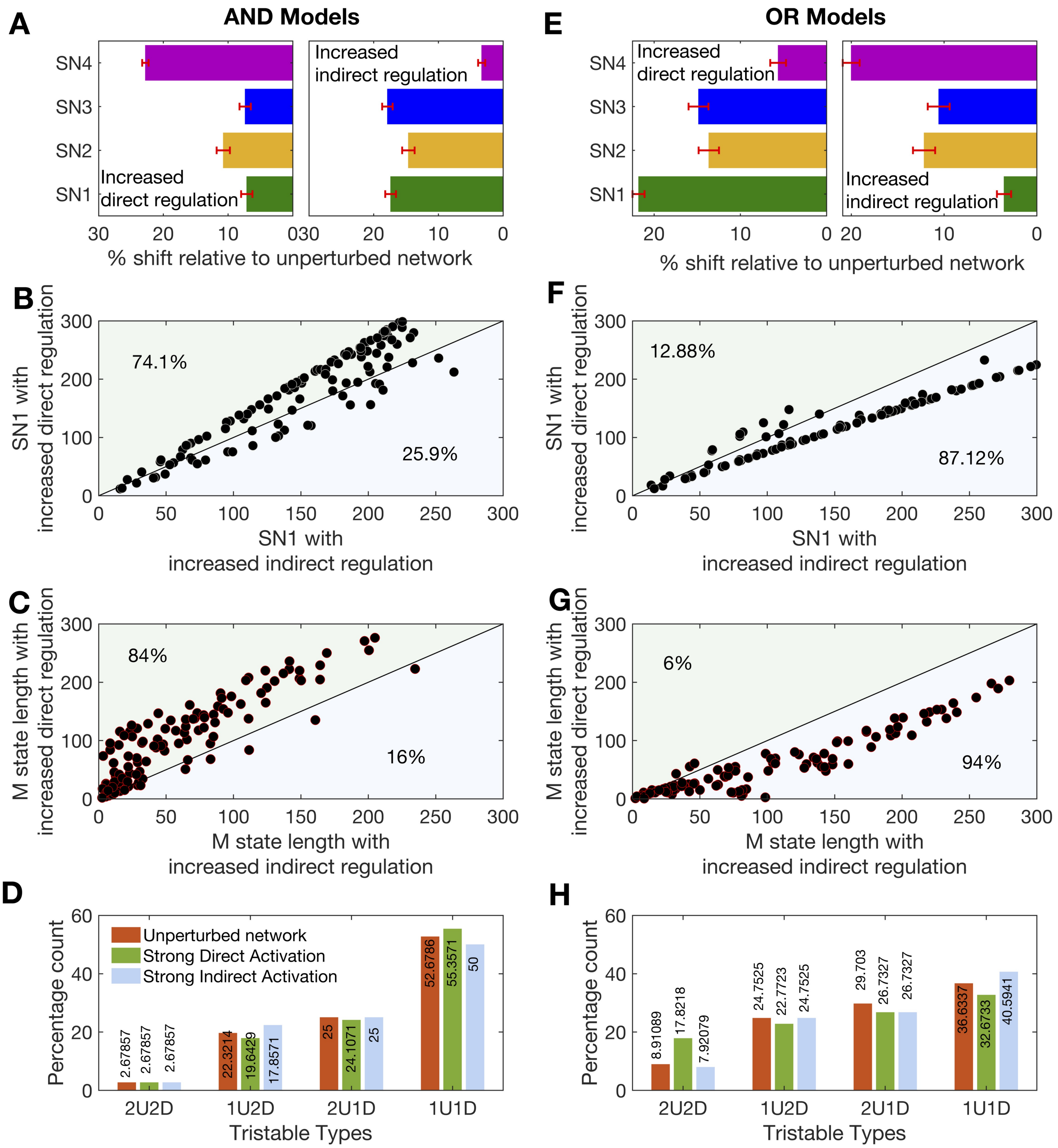


**S3 Figure.** Impact of varying regulation parameters by 25% for unconstrained AND/OR models. **A**. Sensitivity of the SN points for unconstrained tristable AND models perturbed by a 25% increase in the direct or the indirect activation strength. **B**. Scatter plots of SN1 (the EMT initiation point) for 25% perturbations to unconstrained AND models. **C**. Scatter plots of the M state length for 25% perturbations to AND models. **D**. The occurrence of different tristable response types for unperturbed models (red bars), direct activation perturbed by 25% (green bars), and indirect activation perturbed by 25% (blue bars) for AND models. **E**. Sensitivity of the SN points for unconstrained tristable OR models perturbed by a 25% increase in the direct or the indirect activation strength. **F**. Scatter plots of SN1 (the EMT initiation point) for 25% perturbations to OR models. **G**. Scatter plots of the M state length for 25% perturbations to OR models. **H**. The occurrence of different tristable response types for unperturbed models (red bars), direct activation perturbed by 25% (green bars), and indirect activation perturbed by 25% (blue bars) for OR models.

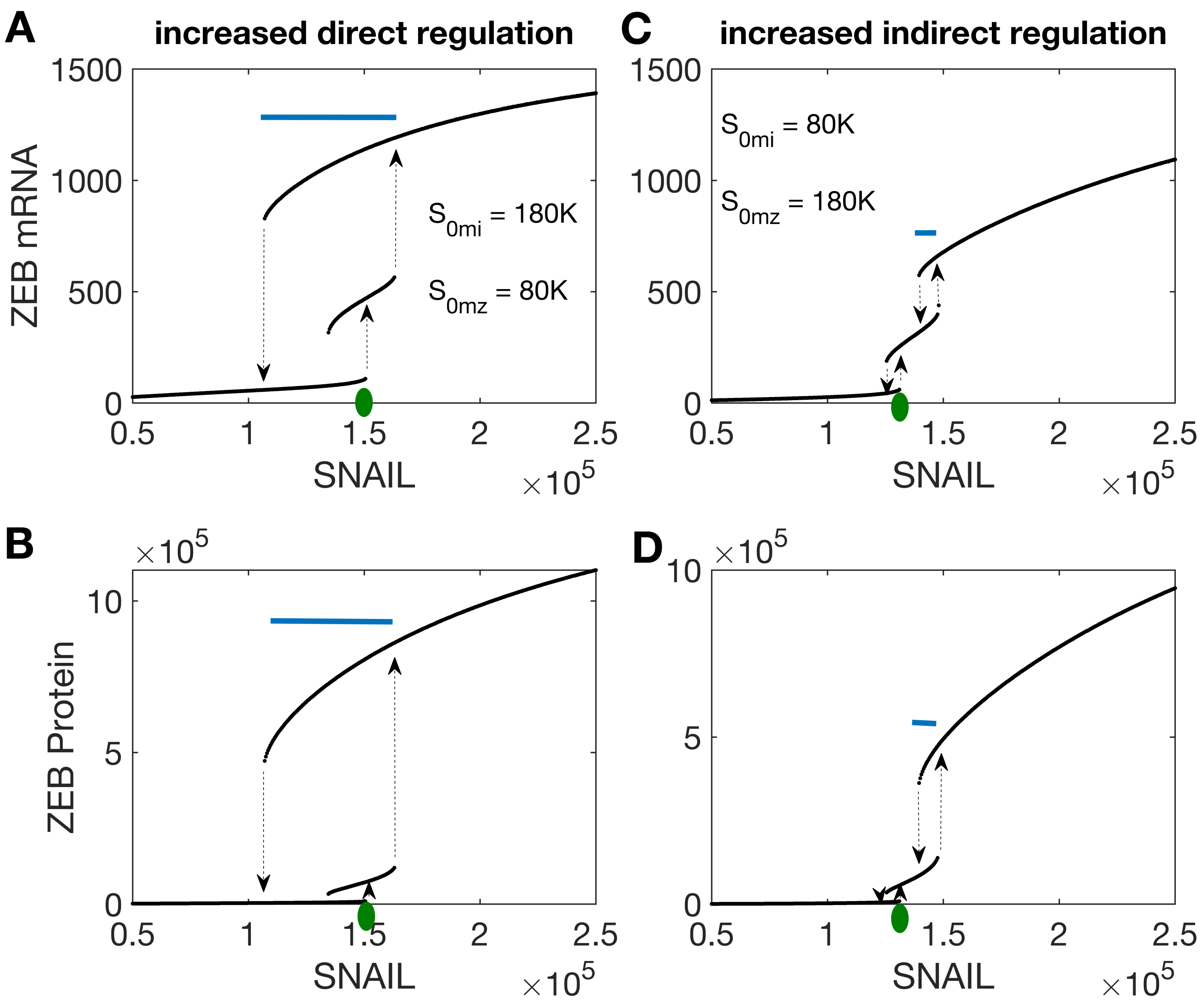


**S4 Figure.** Analysis of the effects of direct vs indirect regulation parameters in the AND logic model of Lu et al. (2013). **A**. ZEB mRNA responses to SNAIL as the direct regulation parameter is increased by 55.5%. Initiation point of EMT (green circle) and length of the M state (blue bar). **B**. ZEB protein responses to SNAIL as the direct regulation parameter is increased by 55.5%. **C**. ZEB mRNA responses to SNAIL as the indirect regulation parameter is increased by 55.5%. **D**. ZEB protein responses to SNAIL as the indirect regulation parameter is increased by 55.5%.

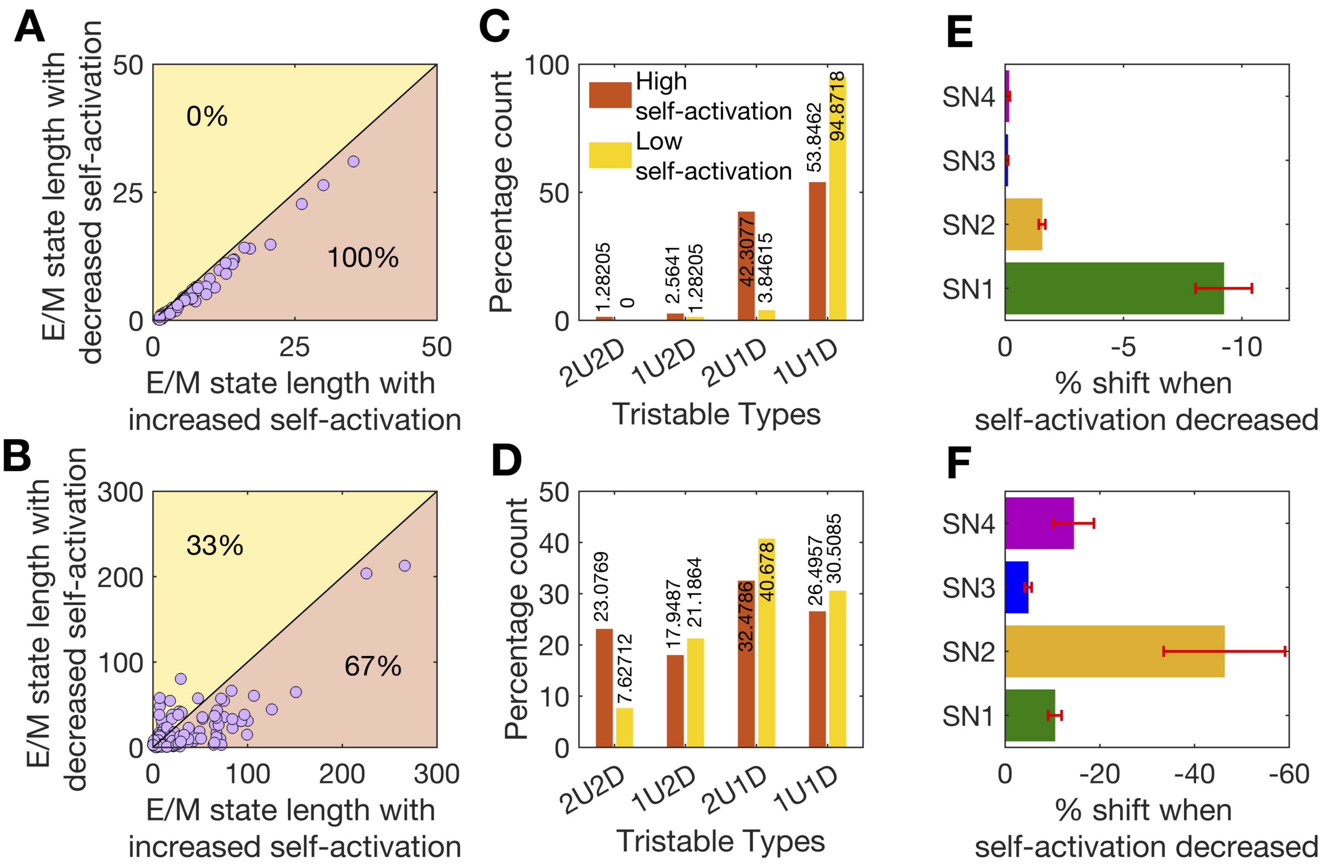


**S5 Figure.** Analysis of the effects of the self-regulation parameter $B_{B0}$. **A-B.** Scatter plot of the hybrid E/M state length for perturbations in parameter $B_{B0}$ for 144 AND logic models (A) and 108 OR logic models (B). **C-D**. The occurrence of different tristable response types for models with self-activation increased (red bars) or self-activation decreased (yellow bars) for 144 AND models (C) and 108 OR models (D). **E-F**. Sensitivity of the SN points perturbed by a 25% decrease in the self-activation strength for AND models (E) or OR models (F).

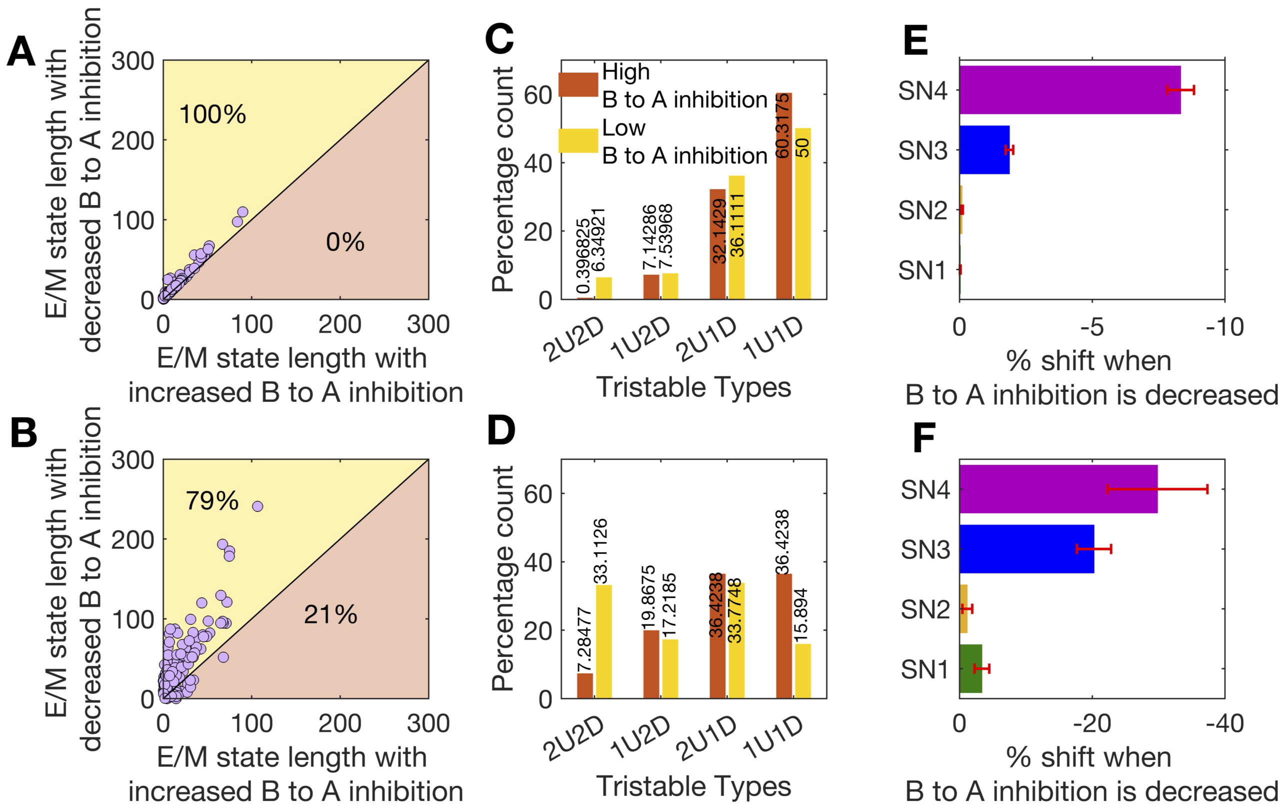


**S6 Figure.** Analysis of the effects of the inhibition parameter $B_{A0}$. **A-B.** Scatter plot of the hybrid E/M state length for perturbations in parameter $B_{B0}$ for 144 AND logic models (A) and 108 OR logic models (B). **C-D**. The occurrence of different tristable response types for models with self-activation increased (red bars) or self-activation decreased (yellow bars) for 144 AND models (C) and 108 OR models (D). **E-F**. Sensitivity of the SN points perturbed by a 25% decrease in the self-activation strength for AND models (E) or OR models (F).

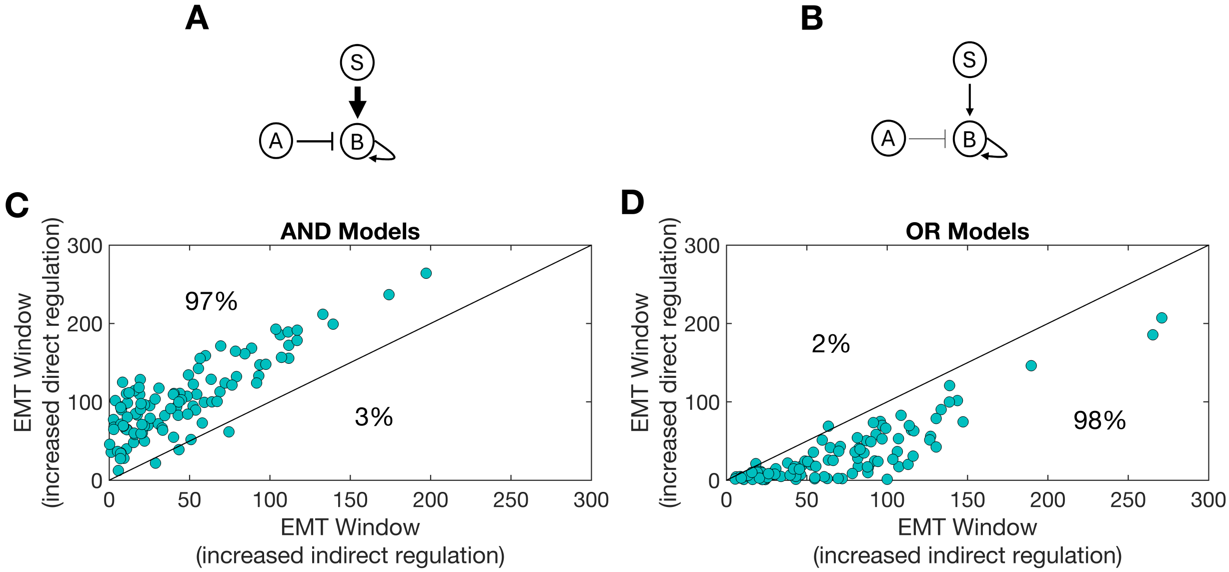


**S7 Figure.** **A**. Scatter plot of the size of the EMT window (distance from SN1 to SN4) for perturbations to the direct vs indirect regulation parameters for 144 AND logic models. **B**. Scatter plot of the size of the EMT window (distance from SN1 to SN4) for perturbations to the direct vs indirect regulation parameters for 108 OR logic models.

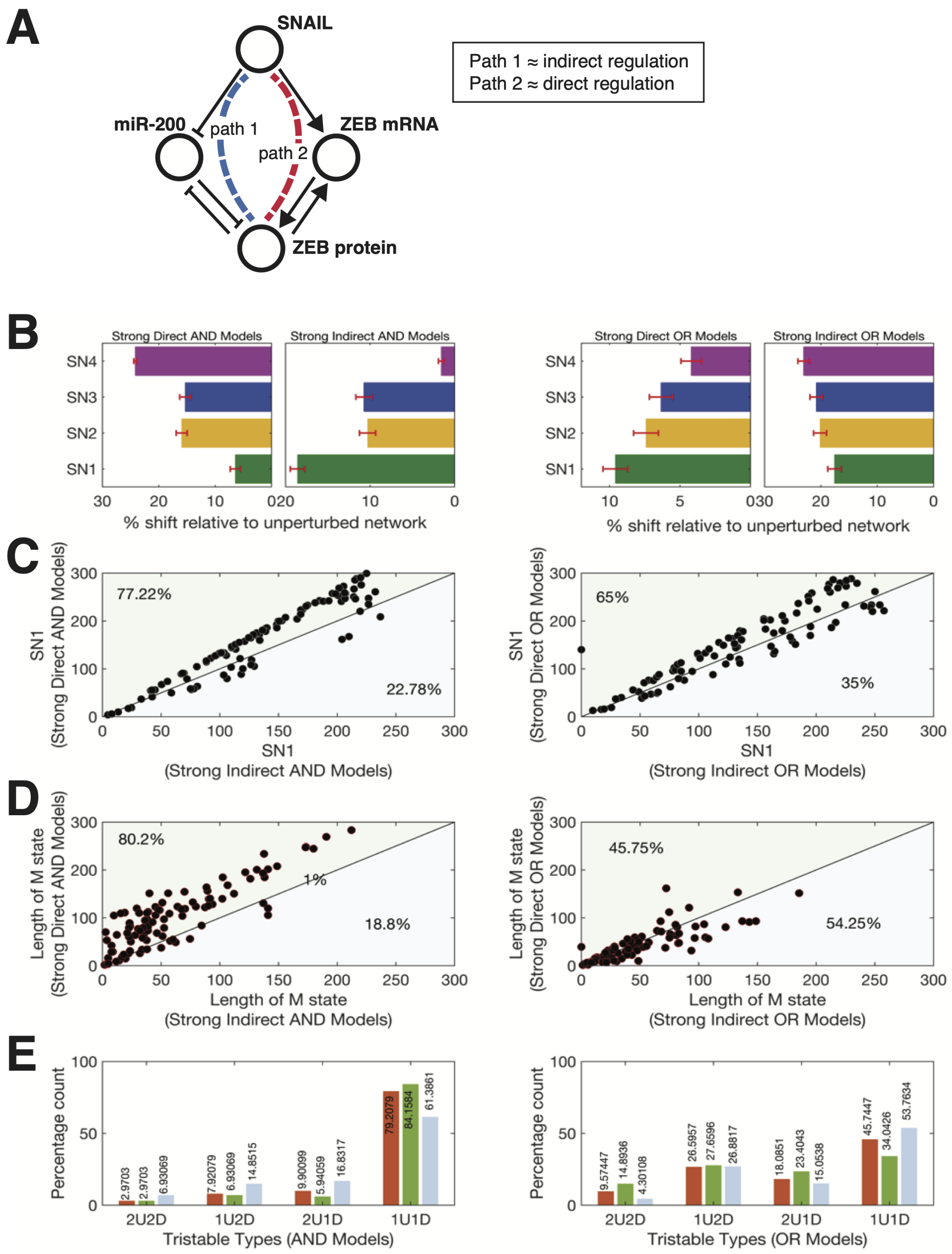


**S8 Figure. Features of EMT landscapes for AND vs OR models incorporating mRNA. A**. Schematic of the EMT network model that incorporates both mRNA and protein species of ZEB. Path 1 is analogous to indirect regulation in previous models; path 2 analogous to direct regulation. **B.** Sensitivity of the SN points for EMT networks perturbed by a 25% increase in the path 2 (indirect) or path 1 (indirect) regulation strength for AND models (left) or OR models (right). **C.** Scatter plots of SN1, the EMT initiation point, for perturbations of the model in (A) constructed with AND logic (left) or OR logic (right). **D**. Scatter plots of the M state length for perturbations of the model constructed with AND logic (left) or OR logic (right). **E.** The occurrence of different tristable response types for unperturbed models (red bars), path 2 perturbed by 25% (green bars), and path 1 perturbed by 25% (blue bars) for AND models (left) and OR models (right).
